# Dynamics of preventive and reactive cancer control using low-impact treatments

**DOI:** 10.1101/014589

**Authors:** Andrei R. Akhmetzhanov, Michael E. Hochberg

**Affiliations:** Institute of Evolutionary Sciences of Montpellier - UMR 5554, University of Montpellier II, CC065, Place Eugéne Bataillon, 34095 Montpellier Cedex 5, France; Theoretical Biology Lab, Dept. of Biology, McMaster University, Hamilton, Ontario L8S4K1, Canada; Santa Fe Institute, Santa Fe, NM 87501, USA; Wissenschaftskolleg zu Berlin, Wallotst. 19, 14193 Berlin, Germany; Kavli Institute for Theoretical Physics, University of California, Santa Barbara, CA 93106-4030, USA

**Keywords:** evolution of resistance, chemoprevention, tumor management

## Abstract

Cancer poses danger because of its unregulated growth, development of resistant subclones, and metastatic spread to vital organs. Although the major transitions in cancer development are increasingly well understood, we lack quantitative theory for how preventive measures and post-excision (’reactive’) treatments are predicted to affect risks of obtaining a life threatening cancer or relapse, respectively. We employ analytical and numerical models to evaluate how continuous measures such as life style changes, and certain non-targeted and targeted treatments affect both neoplastic growth and the frequency of resistant clones. We find that preventive measures can have a negligible impact on pre-cancerous lesions and yet achieve considerable reductions in risk of invasive cancer. Importantly, our model, based on realistic parameter estimates, predicts that daily cancer cell arrest levels of 0.2–0.3% produce optimal outcomes for prevention, whereas for reaction the level is 0.3–0.4%. For similar cancer cell populations, prevention outcomes are, on average, always better than reactive ones. This is because reactive measures are more likely to select for faster growing subclones with higher probabilities of resistance, highlighting the difficulty in countering relapse regardless of therapeutic impact on cancer cell populations. We discuss these results and other important mitigating factors that need to be taken into consideration in a comparative understanding of preventive versus reactive treatments.

## Introduction

Mathematical models play an important role in describing and analyzing the complex process of carcinogenesis. Natural selection for increases in tumor cell population growth can be represented as the net effect of increased fission rates and/or decreased apoptosis (e.g., [1]). Relatively rare driver mutations confer such a net growth advantage, whereas numerically dominant passenger mutations with initially neutral or mildly deleterious effects [2–4] can initially grow in frequency due to genetic hitchhiking or subsequent selection. Amongst the many passengers in a growing tumor, some can contribute to chemoresistance, and sufficiently large tumors could contain different clones that, taken as a group, can resist some, if not most, possible chemotherapies (see [5] for resistance to imatinib). Chemotherapeutic remission followed by relapse suggests that these resistant cells are often present at low frequencies prior to therapy, either due to genetic drift or costs associated with resistance. Resistant phenotypes subsequently increase in frequency during radiotherapy or chemotherapy, and through competitive release, they may incorporate one or more additional drivers, resulting in accelerated growth compared to the original tumor [6].

Previous mathematical studies have considered alternatives to attempting to minimize or eradicate clinically diagnosed cancers with maximum tolerated doses (MTD) of chemotherapeutic drugs. This body of work indicates that MTD is particularly prone to select for chemoresistance (e.g., [7–9], and what little empirical work exists supports this basic prediction [10], but see [11] for other disease systems). Numerous alternatives to the goal of cancer minimization/eradication have been investigated (e.g., [2, 7, 12–14]). For example, Komarova and Wodarz [12] considered how the use of one or multiple drugs could prevent the emergence or curb the growth of chemoresistance. They showed that the evolutionary rate and associated emergence of a diversity of chemoresistant lineages is a major determinant in the success or failure of multiple drugs versus a single one. Lorz and coworkers [9] recently modeled the employment of cytotoxic and cytostatic therapies alone or in combination and showed how combination strategies could be designed to be superior in terms of tumor eradication and managing resistance than either agent used alone. Foo and Michor [7] evaluated how different dosing schedules of a single drug could be used to slow the emergence of resistance given toxicity constraints. One of their main conclusions is that drugs slowing the generation of chemoresistant mutants and subsequent evolution are more likely to be successful than those only increasing cell death rates.

These and other computational approaches have yet to consider the use of preventive measures to reduce cancer-associated morbidity and mortality, whilst controlling resistance. Prevention includes life-style changes and interventions or therapies in the absence of detectable invasive carcinoma (e.g., [15–18]), for example reduced cigarette consumption [23] or chemoprevention [24]. In depth consideration of preventive measures and their likely impact on individual risk and epidemiological trends is important given the virtual certitude that everyone has pre-cancerous lesions, some of which may transform into invasive carcinoma [19, 20], and concerns as to whether technological advances will continue to make significant headway in treating clinically detected cancers [21, 22].

Here we model how continuous, constant measures affect tumor progression and the emergence of resistant lineages. Importantly, we consider cases where the measure may select for the evolution of resistant phenotypes and cases where no resistance is possible. Our approach is to quantify the daily extent to which a growing neoplasm must be arrested to either eradicate a cancer cell population or to delay a potentially lethal cancer. Several authors have previously argued for how constant or intermittent low toxicity therapies either before or after tumor discovery could be an alternative to maximum tolerated dose chemotherapies [18,25], but to our knowledge no study has actually quantified the modalities (treatment start time, dose) for such approaches using empirically derived parameter estimates [2, 26, 27]. Below we employ the terms ‘treatment’, ‘measure’, and ‘therapy’ interchangeably, all indicating intentional measures to arrest cancer cells.

We first derive analytical expressions for the expected total number of cells within a tumor at any given time. We explore dynamics of both tumor sizes at given times, and times to detection for given tumor sizes. Specifically, we show that the expected mean tumor size in a population of subjects can be substantially different from the median, since the former is highly influenced by outliers due to tumors of extremely large size. We then consider constant daily preventive measures, and show that treatment outcome is sensitive to initial conditions, particularly for intermediate sized tumors. Importantly, we provide approximate conditions for tumor control both analytically and corresponding impact estimates based on our empirical parameter estimates. Finally, we apply our model to reactive therapies following tumor discovery and excision, showing, as expected, that higher impacts on the cancer cell population is required to achieve a level of control comparable to prevention.

## Modeling framework

Previous study has evaluated the effects of deterministic and stochastic processes on tumor growth and the acquisition of chemoresistance ([12, 27, 28], see review [29]). We first consider both processes through exact solutions and numerical simulations of master equations, using the mean field approach. A mean field approach assumes a large initial number of cells [30] and averages any effects of stochasticity, so that an intermediate state of the system is described by a set of ordinary differential equations (i.e., master equations; [31]). Solutions to these are complex even in the absence of the explicit consideration of both drivers and passengers [33, 34].

We do not explicitly model different pre-cancerous or invasive carcinoma states. Rather, our approach follows the dynamics of the relative frequencies of subclones, each composed of identical cells [35,36]. We simulate tumor growth using a discrete time branching process for cell division [27,32,37]. For each numerical experiment, we initiate a tumor of a given size and proportion of cells resistant to the measure under consideration within a tumor.

Briefly, the model framework is as follows. Each cell in a population is described by two characteristics. The first is its resistance status to the measure, which is either “not resistant” (*j* = 0) or “resistant” (*j* = 1). The second property is the number of accumulated driver mutations (maximum *N*) in a given cell line. At each time step cells either divide or die, and when a cell divides, its daughter cell has a probability *u* of producing a driver mutation and *v* of producing a resistant mutation. We assume no back mutation and that cells do not compete for space or limiting resources (see Discussion).

The fitness function *f_ij_*, the difference between the birth and death rates of a cell, is defined by the number of accumulated drivers (*i* = 0, 1,…, *N*) and resistance status (*j* = 0, 1): a sensitive cancerous cell with a single driver has selective advantage *s*, and any accumulated driver adds *s* to fitness, while resistance is associated with a constant cost *c*. Exposure to a single treatment affects only non-resistant cells (*j* = 0), incurring a loss σ to their fitness. Thus, the fitness function is:

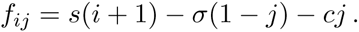

The assumption of driver additivity is a special case of multiplicative fitness, and both are approximately equivalent for very small *s*.

We conducted numerical experiments each with the same initial states, but each using a unique set of randomly generated numbers of a branching process. For each simulation and each time step, the number of cells at time (*t*+1) was sampled from a multinomial distribution of cells at time *t* (see [27,37] for details). Other methodological details can be found in Supplementary information.

Table 1 presents baseline parameter values employed in this study.

**Table 1.** Baseline parameter values.

| Parameter | Variable | Value | Reference |
| --- | --- | --- | --- |
| Time step (cell cycle length) | $T$ | 4 days | [27] |
| Selective advantage | $s$ | 0.4% | [27] |
| Cost of resistance | $c$ | 0.1% | |
| Mutation rate to acquire an additional driver | $u$ | $3.4 \times 10^{-5}$ | [27] |
| Mutation rate to acquire resistance | $v$ | $10^{-6}$ | [12] |
| Maximal number of drivers | $N$ | 9 | |
| Initial cell population | $n(0)$ | $10^6$ cells | |
| Pre-resistance level | $\kappa$ | 0.01% | [32] |
| Number of replicate numerical simulations<br>(excl. the ones with extinction) | - | $10^6$ | |
| Detection threshold | $M$ | $10^9$ cells | [62] |

## Results

### Preventive Measures

We first study mean-field dynamics by considering the distribution of tumor sizes at different times and examine effects on the mean (see Supplementary information). Numerical experiments were carried out by assuming that tumors contained 10^6^ cells when treatment commenced and, importantly, had neither additional drivers nor resistance mutations (*i* = 0, *j* = 0). This is obviously an oversimplification, and we relax these assumptions below and in the next section.

Figure 1**A** shows the excellent correspondence between numerical experiments and analytical results for *σ* on the order of *s*. A more detailed study of the distribution of tumor sizes reveals that the mean *n*(*t*) diverges considerably from median behavior in the majority of cases, since the former is strongly influenced by outliers with high tumor cell numbers (see Figure 1**B** and Supplementary figure 1).

**Figure 1.**
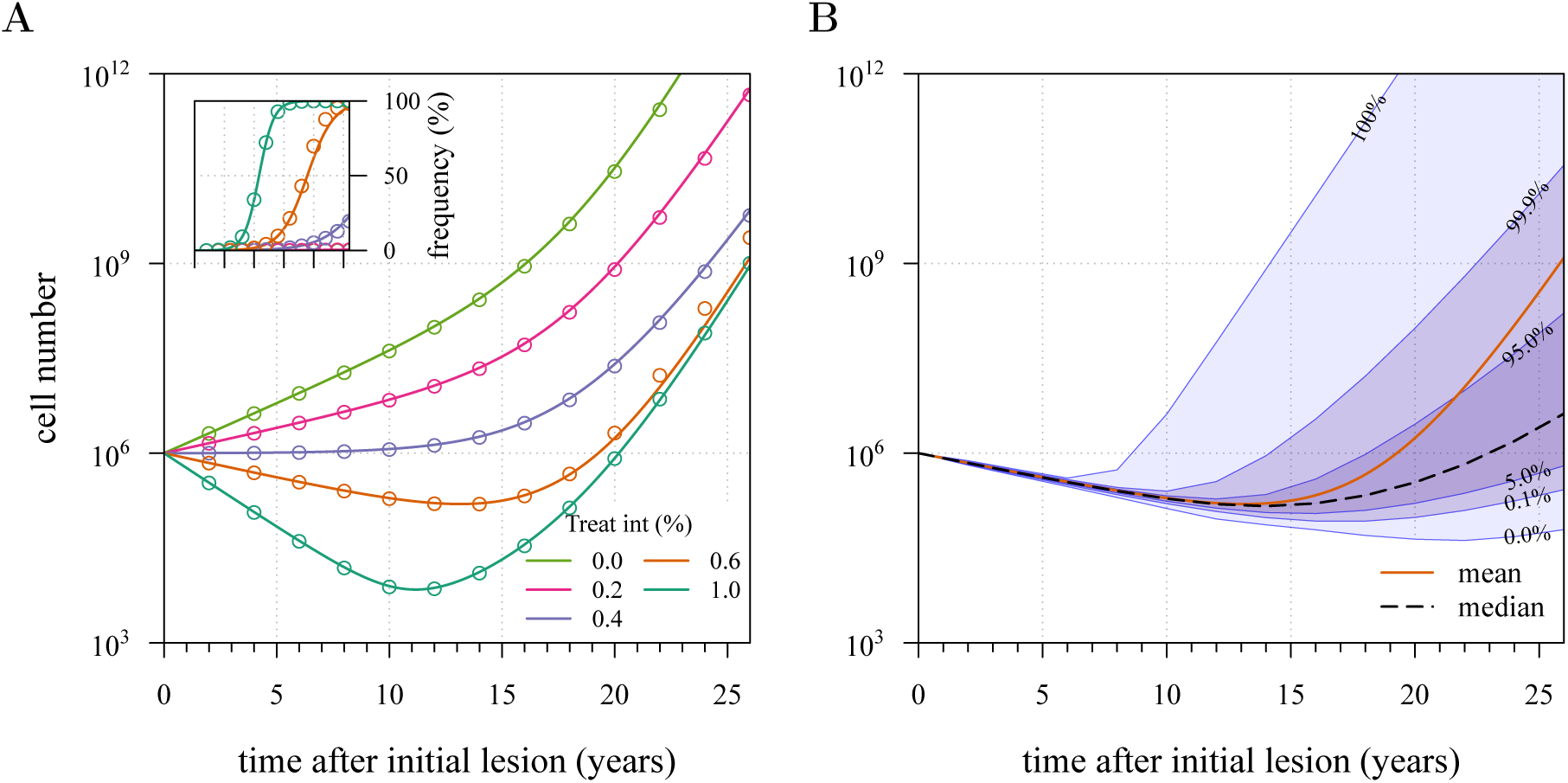
Mean field dynamics concord with numerical simulations. (**A**) Effect of treatment level and observation time on mean tumor size. (**Inset**) Mean frequency of resistant cells within tumors corresponding to three of the cases in **A**. Lines are analytically computed mean-field trajectories, while dots are numerical simulations (see Supplementary information for details). (**B**) Dynamics of mean and median tumor size, and percentiles around the mean (shaded areas), assuming a fixed constant treatment of *σ* = 0.6%. Treatments start at *t* = 0, and the maximal number of additionally accumulated drivers is 3. See Table 1 for other parameter values.

Figure 2 shows four examples of numerical experiments. An untreated tumor reaches the detection threshold of 10^9^ cells by *c*. 18 years on average, and because it is not subject to strong negative selection (we assume low *c*), any emerging resistant cell-lines remain at low frequency (0.03% at the time of detection in the example of Figure 2**A**). In Figure 2**B**, low treatment intensity delays tumor growth and thus time of detection by *c*. 16 years, while an increase in dose tends to result in tumors dominated by resistant cells (Figure 2**C**). Despite being unaffected by treatment, resistant cell populations sometimes observed to go extinct due to stochasticity (Figure 2**D**), and this tends to occur more at high treatment levels, because there are fewer sensitive tumor cells to seed new (mutant) resistant cell populations.

**Figure 2.**
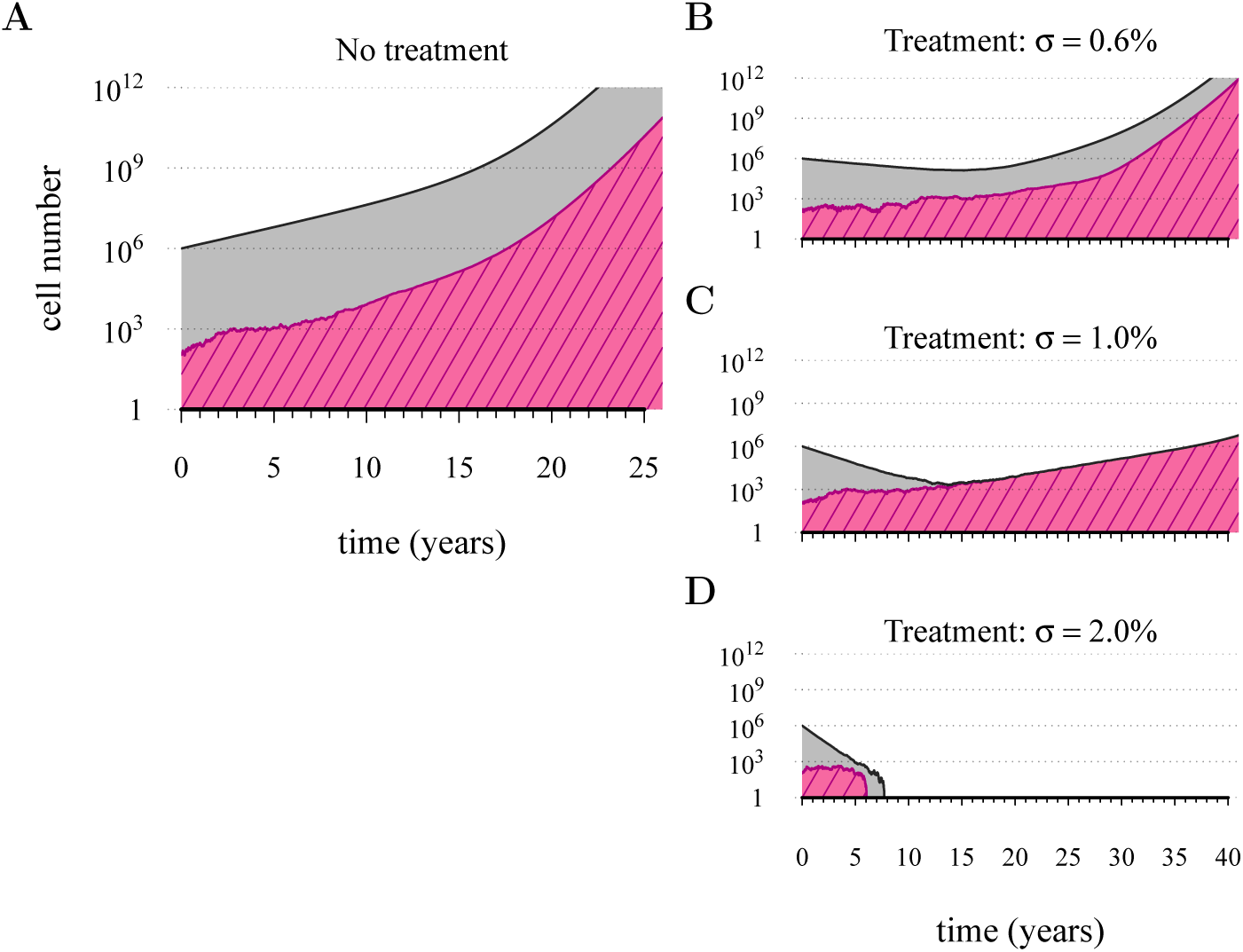
Treatments curb or eliminate tumors. Examples of single patient tumor growth for (**A**) no treatment; (**B**) *σ* = 0.6%; (**C**) *σ* = 1.0%; (**D**) *σ* = 2.0%. The shaded area shows the change in total tumor size, and the hatched area the resistant part of a tumor. Parameter values are as in Table 1.

We next considered how therapies affected the distribution of tumor detection times in cases where the cancer cell population attained the threshold of 10^9^ cells. The magnitude of the selective advantage *s* shows that tumor growth is largely driven by its non-resistant part for relatively low impact treatments *σ* < 2*s* (Figure 3). Importantly, the tumor shifts from being mainly non-resistant to resistant at *σ* ≈ 2*s*, which is reflected by the inflection point in the trajectory of the median (indicated by point *B* in Figure 3). Notice that detection times are also most variable at *σ* ≈ 2*s*. The median changes smoothly at high treatment levels (*σ* > 2*s*), tending to a horizontal asymptote. This is explained by the fact that the sensitive part is heavily suppressed at high treatment levels, meaning that the dynamics are strongly influenced by the actual time point at which the resistance mutation occurs.

**Figure 3.**
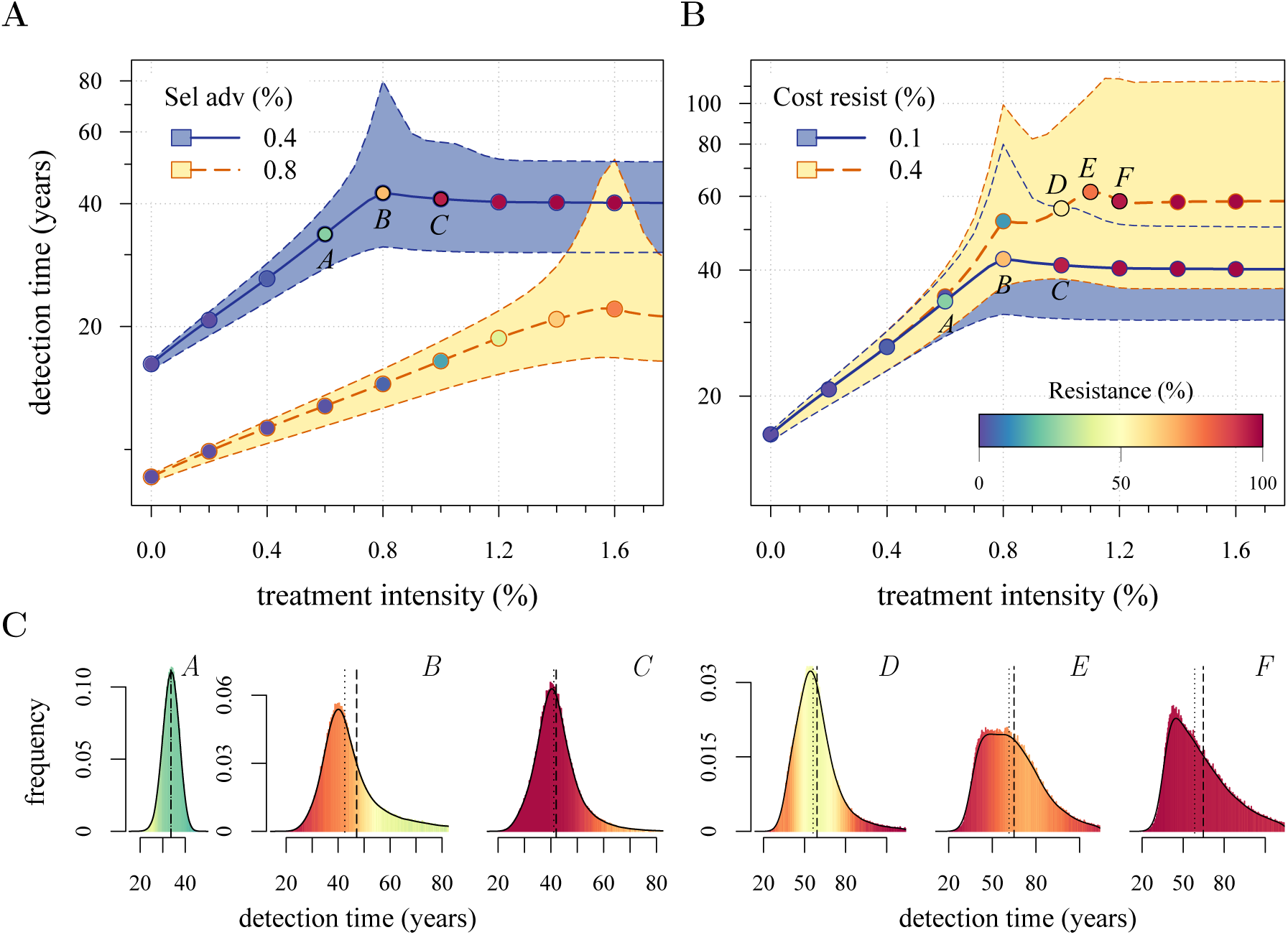
Treatment level affects both detection time and frequency of resistance. The median and 90% confidence intervals (shaded areas) of detection times, measured as years beyond the initiation of the preventive measure. Effects of: (**A**) the selective advantage and (**B**) the cost of resistance. (**C**) Particular samples of the distribution of detection times for corresponding points, indicated in **A** and **B**. Dashed black line is the mean and the dotted line is the median. Bottom panel shows the mean number of additionally accumulated drivers within tumors over periods of 3 months. Color-code indicates the level of resistance in detected tumors over 3 month intervals (see inset in **B**). All cells *j* = 0 at *t* = 0. Other parameters are as in Table 1. Note that the detection time is log-transformed in **A** and **B**. The treatment intensity *σ* in this an all other figures is measured per cell cycle.

We find, counterintuitively, that early-detected tumors are more likely to be resistant under constant treatments than those detected at later times (*A*, *B*, *C* in Figure 3**C**). This is because tumors under treatment that by chance obtain resistance early grow faster than those that do not. We find that by the time of detection, non-resistant tumors usually accumulate up to 4 additional drivers on average, while resistant tumors have fewer. For larger values of *c*, an additional non-regularity emerges (segment *DEF* in Figure 3**B**), appearing at *σ* ≈ 3*s* and is associated with tumors having a majority of cells with maximum numbers of drivers. This region is also characterized by a different transition to complete resistance (compare Supplementary videos S1 and S2 for relatively low and high costs of resistance, respectively). For example, at point *D* tumors with a majority of non-resistance have less variable detection times than tumors with a majority of resistant cells (**B** and corresponding panel **C** in Figure 3). Treatment levels along the segment *DEF* result in tumors that are more likely to be resistant as one goes from the center to the tails of the distribution of detection times. This differs qualitatively from the previous case of low cost of resistance, where the tumors are less resistant in the tail of the distribution of detection times (Figure 3**C**).

The inflection point at *σ* ≈ 2*s* in Figure 3A is due to the accumulation of additional drivers within tumors and associated increases the likelihood that the tumor eventually resists treatment. Since the initial population consists of 10^6^ cells, in the absence of treatment, a mutant cell with one additional driver and associated fitness 2*s* will appear very rapidly. Such a tumor can be suppressed only if we apply the treatment with *σ* > 2*s*. This is supported by additional numerical experiments, where we vary the maximal number of additional driver mutations *N*: the inflection point *σ*≈2*s* disappears when *N* = 0 (Figure 4**A**). The inflection points at *σ* = 3*s*, 4*s* emerge at treatment levels that effectively suppress sensitive subclones with the most drivers before resistance mutations are obtained (*cf* Figures 4**A**,**C**–**D** with Figure 4**B** and Supplementary Video S3). Specifically, the peaked distributions, corresponding to better therapeutic outcomes, tend to disappear when resistant subclones emerge.

**Figure 4.**
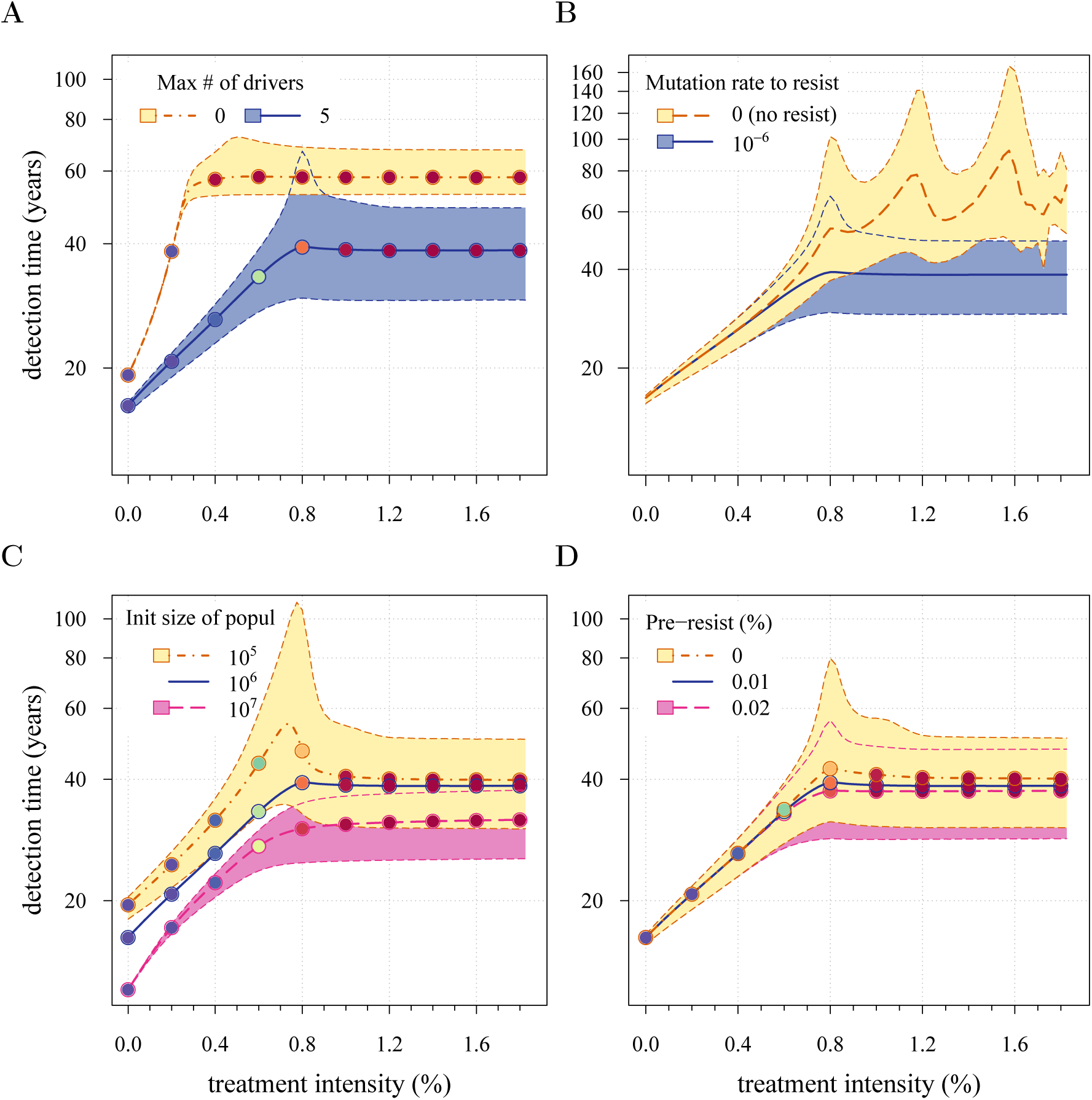
Sensitivity analysis of several key parameters. The median (thick line) and 90% confidence intervals (shaded areas with dashed boundaries) for the distribution of detection times. Parameter values are as in Table 1 except the one being varied: (**A**) maximal number of additionally accumulated drivers; (**B**) presence of resistant cell-lines; (**C**) initial cell number; (**D**) level of initial partial resistance of a tumor. The color code for points indicates the average level of resistance within tumors, analogous to Figure 3. For simplicity, only the median is indicated in **C** and **D** for the baseline case (blue line).

The initial cell number *n*(0) affects both the median and distribution of detection times (Figure 4**C**). For large initial tumors, growth is deterministic and exponential. As the initial size is decreased from 10^6^ to 10^5^, stochastic effects are increasingly manifested by greater variability in tumor inhibition and an inflection point observed at the 95^th^ percentile. Moreover, we find that a tumor is likely to be eradicated under a range of constant treatments when *n*(0) = 10^5^ or fewer initial cells; in contrast, a tumor is virtually certain to persist regardless of treatment level for *n*(0) = 10^7^ cells or greater (Supplementary figure 2**A**, **B**). In other words, our model indicates that tumors that are *c*. 1% the size of most clinically detectable, internal cancers will typically be impossible to eradicate.

The above analysis assumes zero initial resistance within a tumor. Given the mutation rates assumed here, we can expect that many tumors with one million cells will already contain resistant cells (38). As shown in Figure 4**D** larger values of existing resistance create a transition from stochastic to deterministic tumor growth, and worse control outcomes (Supplementary figure 2**C**).

### Reactive measures

We numerically investigated measures starting after tumor discovery and excision, by initiating a primary tumor with one cell (*i* = 0, *j* = 0), thereafter growing to either 10^9^ (early detection) or 10^11^ (very late detection). At detection we assume that the primary tumor is removed, leaving a small number of residual and/or metastatic cells (10^4^ or 10^6^). We study how a measure (usually some form of chemothrerapy or radiation therapy, but could also involve adjuvants after an initial therapy) after excision affects the probability of treatment success, the distribution of times for tumor relapse and resistance levels. We contrast these resection scenarios with prevention, and assume that the undetected tumor (*i.e.*, the target of prevention) grows from a single cell and accumulates drivers and resistance mutations. (Recall that in the previous section we assumed that when a measure commenced, preventively treated tumors had neither additional drivers nor resistance mutations (*i* = 0, *j* = 0)). Distributions of driver mutations for each scenario are presented in Figure 5**A** and Supplementary figure 3**A**.

**Figure 5.**
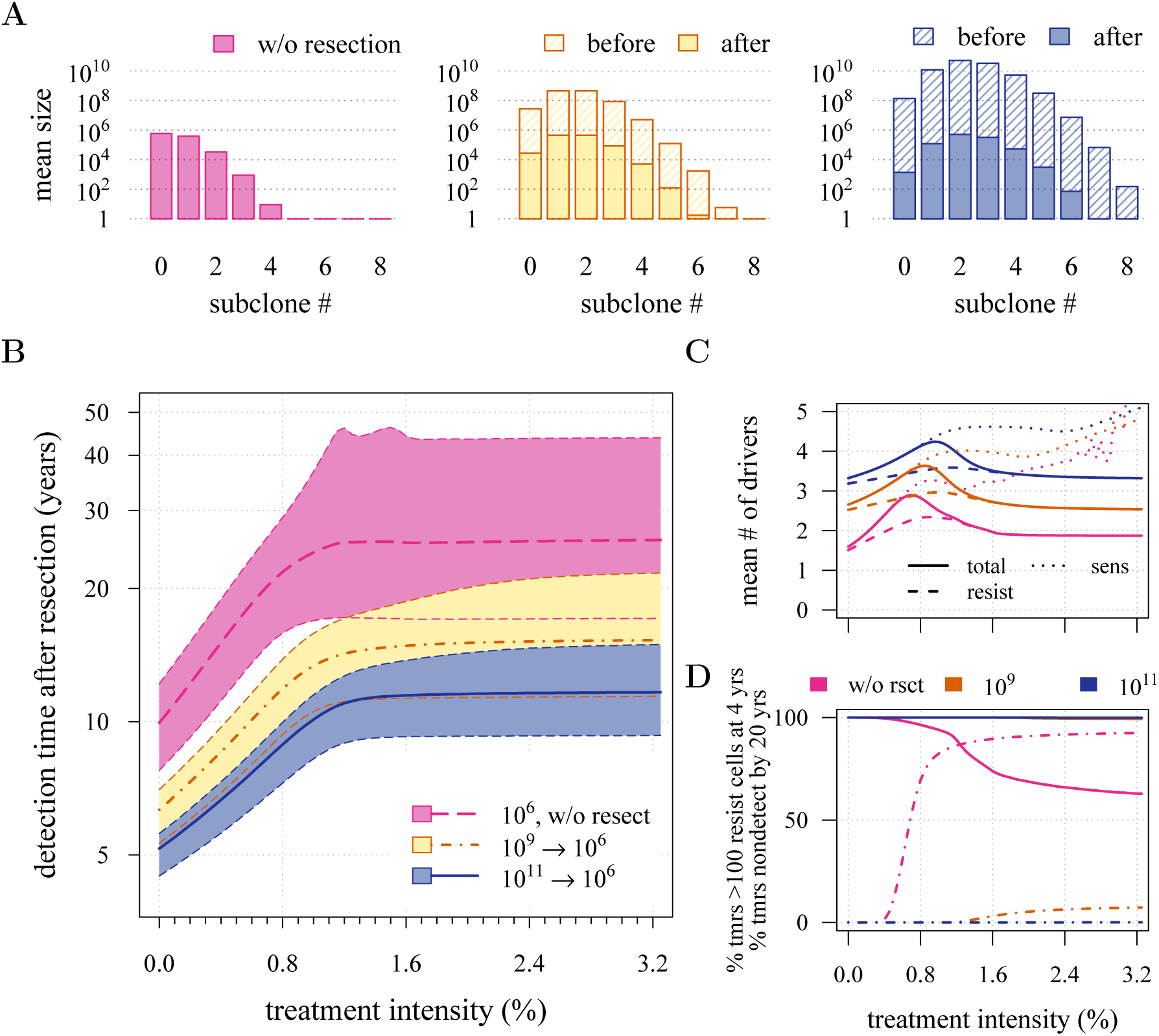
Effects of preventive and reactive measures. Therapeutic outcomes when 10^6^ cancer cells remain after tumor resection. (**A**) The distribution of mean sizes of subclones (hatched bars = before removal and solid bars = post removal). (**B**) The distribution of detection times (thick line = median, filled area with dashed boundaries = 90% CIs) for different constant treatment intensities. (**C**) The mean number of accumulated drivers within a tumor at the time of detection. (**D**) The percentage of cases when a tumor consists of less than 100 resistant cells at 4 years post-resection (solid lines) and the percentage of cases when tumor sizes are below the detection threshold (dashed-and-dotted lines). Maximal number of additional drivers is 9, other parameter values as the same as in Table 1.

First, we examine the case where 10^6^ cells remain after resection. As suggested by our studies above on prevention, one million cells have a very high probability of already containing, or rapidly obtaining, resistant subclones and deterministic effects dominate subsequent dynamics of tumor growth. Comparing the median expectations of years from tumor excision to relapse, early discovery (at 10^9^ cells) yields an additional 3.4 years compared to late discovery (at 10^11^ cells) at *σ* = 1.5% (medians for low *vs* high detection thresholds are 14.8 and 11.4 years, respectively; Figure 5**B**). Consider the following example. 20 years after resection and commencing treatment, the probability of tumor non-detection (i.e., the tumor is either eradicated or does not reach the detection threshold) is close to zero, regardless of treatment intensity (Figure 5**D**). Contrast this with prevention starting at the same tumor size (10^6^ cells): the detection of potentially life-threatening tumors is substantially later than either of the excision cases (median 25.5 years for *σ* = 1.5%), and tumors are managed below the detection threshold after 20 years occurs in more than 80% of cases for any *σ* > 1.0% (Figure 5**D**).

Now consider a residual population of 1/100^th^ the previous case, that is 10^4^ cells. Here, stochastic effects play a more important role in dynamics (Supplementary figure 3). Due to initial heterogeneity (*i.e.*, the co-occurrence of many subclones), when there are 4 and 5 (5 and 6) additional drivers in the dominant subclones of a residual cancer from an excised tumor of 10^9^ (10^11^) cells, we observe a double peak at 4*s* and 5*s* (5*s* and 6*s*) (*cf* Supplementary figure 3**B**). These peaks in variability of outcomes are a result of the stochastic nature of the appearance of the first resistance mutations and of additional driver mutations. Counterintuitively, the secondary detection times are more variable for small initial tumors compared to larger ones (*cf* the median 35.8 years, 90% CIs [17.0, 70.5] years *vs* 22.4, [13.7, 37.0] years for 10^9^ *vs* 10^11^, respectively, with *σ* = 1.5%). This effect is due to resistance emergence in more aggressive subclones for larger tumors, such that the tumor relapses more deterministically (i.e., with less variability and faster on average). The distribution of the mean number of accumulated drivers within tumors and the probability of tumor non-detection after 20 years are shown in Supplementary figure 3**C** and 3**D**, respectively.

Importantly, for both thresholds of tumor excision, subsequent treatment levels beyond *c. σ* = 1.5% make little difference in terms of tumor growth (Figure 5**D**), since virtually all of the sensitive cells post-excision will be arrested or killed by the measure beyond this level, leaving uncontrollable resistant cells to grow and repopulate the primary tumor site and/or metastases. Moreover we find that for reactive measures, data on time to clinical discovery (a proxy for age at discovery) and driver number are less predictive of therapeutic outcome as treatment intensity increases (Supplementary figure 4–7). We see in particular that knowledge about the number of drivers at the time of tumor discovery is a better predictor of therapeutic outcome than information about the time from tumor initiation to discovery (*cf* Supplementary figure 4, 5 and 6, 7).

The above results consider prevention and reaction as independent rather than alternative approaches. Thus, although prevention delays tumor growth for longer times on average than does resection followed by treatment, because prevention is always initiated before resection, the actual time gained (in terms of age rather than time) by the former versus the latter will be less than the differences reported in Figure 5B and Supplementary figure 3**B**. Figure 6 presents a comparative scenario of prevention versus resection. Prevention may either succeed without recurrence, or should the measure initially fail and a tumor be clinically detected, the patient has a second chance, whereby the tumor is resected and treatment continued (assumed at the same *σ*), either to a further relapse (failure) or non-detection and success (Figure 6**A**). Compare this scenario with the more standard resection followed by treatment, which either results in relapse (and failure), or detection-free life (Figure 6**B**).

**Figure 6.**
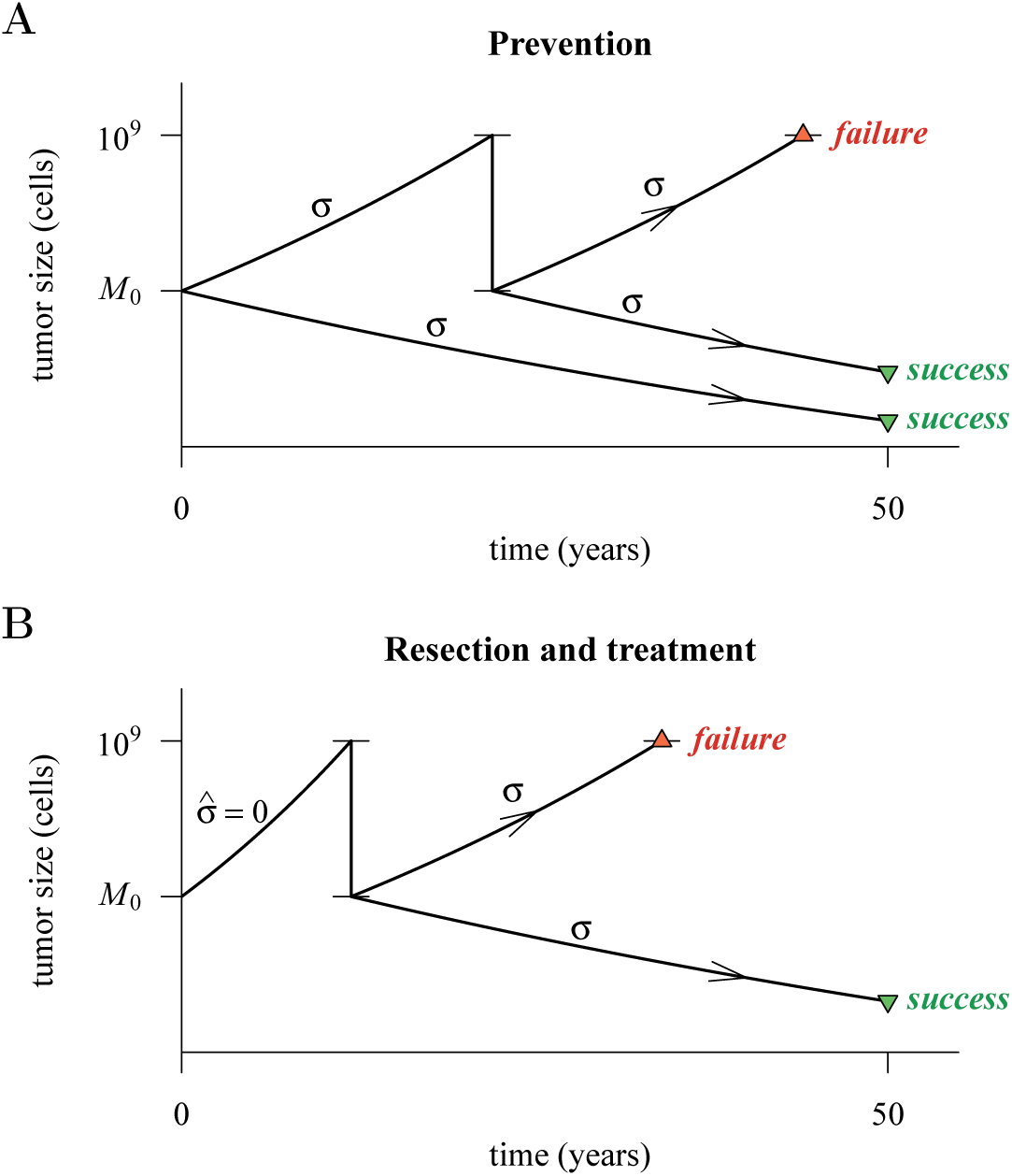
Hypothetical process of preventive and reactive measures. A tumor is initiated by one cell, and grows to size *M*_0_ (10^4^ or 10^6^ cells). It is then either treated constantly with intensity *σ* (Prevention: **A**) or continues to grow 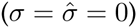 to 10^9^ cells (Reaction: **B**), whereupon it is resected to *M* = *M*_0_ cells and then treated with intensity *σ*. A treatment fails should the tumor attain 10^9^ cells a second time, by 50 years after the initial lesion of size *M*_0_; otherwise the treatment is deemed a success.

Figure 7 presents the comparative outcomes. We see that when prevention starts at (or reactive resection results in) relatively large cancer cell populations (one million cells), prevention is superior to reaction, should relapse occur (Figure 7**A**), but only results in small comparative gains in outright treatment success (Figure 7**E**). Resected tumors in both the prevention and reaction scenarios contain abundant resistant cells (Figure 7**C**). In contrast, when prevention is started very early or the efficiency of resection is high, such that both events are associated with lower cancer cell populations (10000 cells), both prevention and reaction equally delay cases of relapse (Figure 7**B**). Because the number of residual cells following resection is much lower than the first scenario, some resected tumors in the sample will be initially resistance free (Figure 7**D**). This, together with the smaller residual population and fewer subclones in the highest driver classes contribute to improved outcomes should relapse occur (Figure 7**B**) and overall treatment success at sufficiently high intensities (Figure 7**F**).

**Figure 7.**
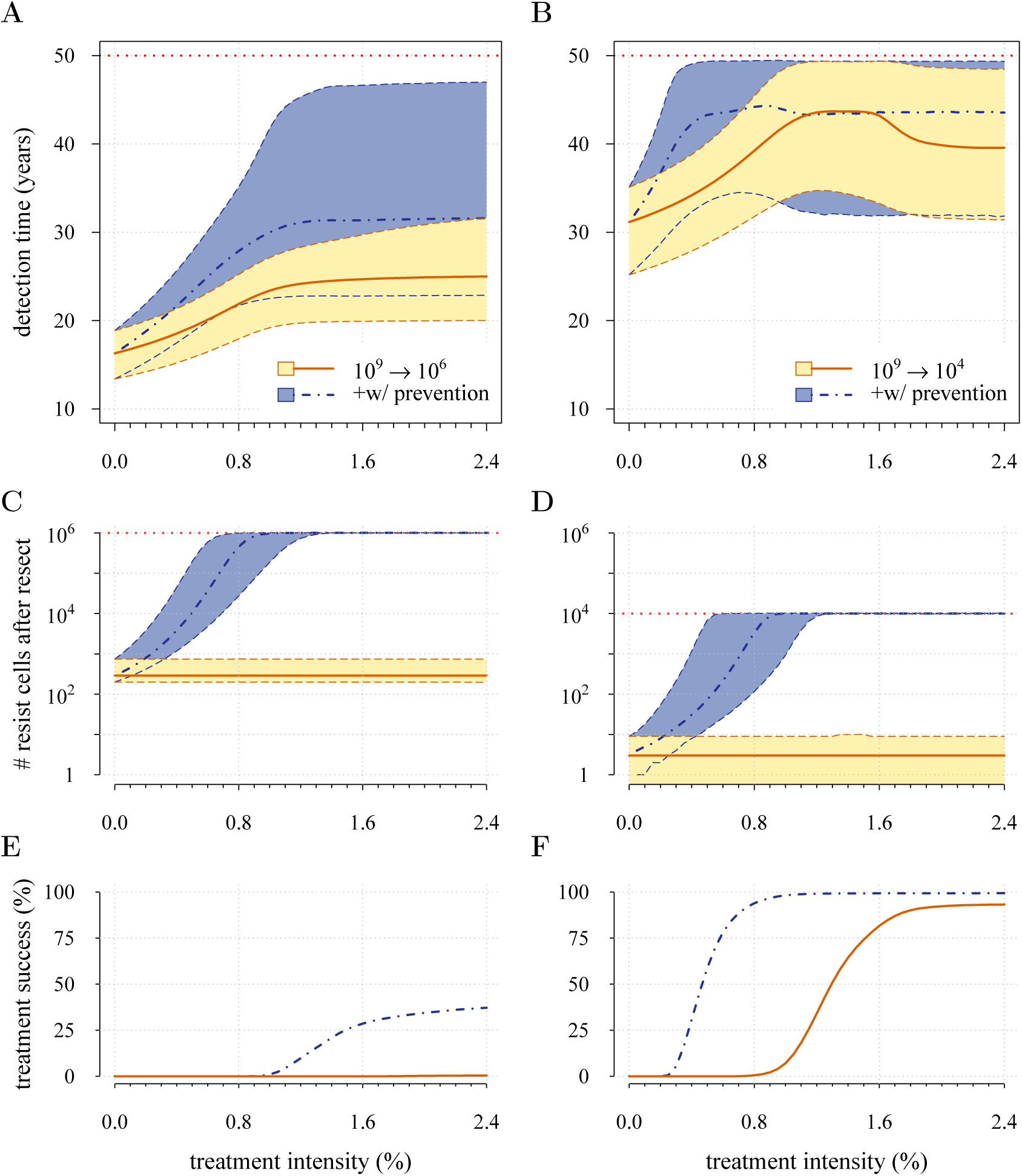
Comparison of preventive and reactive strategies. Tumors are either treated at *M*_0_ = 10^6^ cells (left panels) or *M*_0_ = 10^4^ cells (right panels). Prevention (blue lines and shading) versus reaction (red lines and yellow shading). (**A**, **B**) Distribution of times to relapse for treatment failures. (**C**, **D**) Resistant cell populations after initial failure (red line and yellow shading = population just after resection). (**E**, **F**) Probability of treatment success, defined as the proportion of cases where the tumor remains undetected (either extinct or below 10^9^ cells) by 50 years after the initial lesion. Parameters as in Table 1. See Figure 5 for details.

## Discussion

Maximum tolerated dose chemotherapies present numerous challenges, a major one being the selection of resistant phenotypes, which are possible precursors for relapse [39]. We mathematically investigated how the intensity of an anti-cancer measure, modeled as the arresting effect on a cancer cell population, resulted in success (i.e., either eradication or long-term tumor control) or failure (tumors growing beyond a threshold indicative of a life threatening cancer). Our central result is that beyond low impact thresholds, no additional control is achieved, and resistant subclones are selected. We find that maximal control occurs at surprisingly low daily levels of arrest: approximately 0.2% and 0.3% for preventive and reactive therapies, respectively. An important result of our study is that prevention almost invariably results in greater treatment success than comparable preventive control (Figures 5,7).

We considered two contrasting scenarios. In the first, people at high risk of contracting a life threatening cancer make life-style changes or receive continuous, chemopreventive therapies, and in the second, more usual situation, a tumor is discovered, removed and the patient treated with specific cytotoxic or cytostatic chemicals and/or with radiation. We found that, as expected, prevention requires smaller effects on tumor populations of a given size than do measures post-excision, the latter having smaller probabilities of complete cure and shorter times to tumor relapse. That resistant cell lines accumulate drivers through time suggests that second-line measures following relapse are less likely to be successful in excision scenarios compared to preventive ones. (This is despite residual cancer cell populations that are likely to manifest higher frequencies of resistance in preventive scenarios (Figures 7**C**,**D**).) This prediction depends on the untested, but reasonable assumptions that multi-driver resistant cell populations grow faster and are more genomically unstable than early primary tumors, and that the former thus accumulate a larger number and diversity of resistant mutations. Our results suggest that adopting slightly lower treatment levels than optimal for delaying tumor growth would be an important strategy to manage resistance, for example, if the resistant phenotype were refractory to multiple drugs [40]. Below we discuss challenges to cancer management for both preventive and reactive scenarios.

### Preventive approaches

Whereas primary prevention is becoming an increasingly significant approach to reducing risk of certain cancers such as breast cancers [41], chemopreventive therapies more generally are uncommon, despite empirical support for their effects [15]. Several theoretical and in vitro experimental studies support the potential for chemoprevention to reduce risks of life threatening cancers. For example, Silva and colleagues [42] parameterized computational models to show how low doses of verapamil and 2-deoxyglucose could be administered adaptively to promote longer tumor progression times. These drugs are thought to increase the costs of resistance and the competitive impacts of sensitive on resistant cancer cell subpopulations. However, some of the most promising results have come from studies employing non-steroidal anti-inflammatory drugs (NSAIDs), including experiments [43], investigations of their molecular effects [44, 45], and their use [46]. For example, Ibrahim and coworkers [43] studied the action of NSAIDs and specifically sodium bicarbonate in reducing prostate tumors in male TRAMP mice (i.e. an animal model of transgenic adenocarcinoma of the mouse prostate). They showed that mice commencing the treatment at 4 weeks of age had significantly smaller tumor masses, and that more survived to the end of the experiment than either the controls or those mice commencing the treatment at an older age. Kostadinov and colleagues [44] showed how NSAID use in a sample of people with Barrett’s esophagus is associated with reductions in somatic genomic abnormalities and their growth to detectable levels. It is noteworthy that it is not known to what extent reductions in cancer progression under NSAIDs is due to either cytotoxic or cytostatic effects, or both. Although we do not explicitly model cytotoxic or cytostatic impacts, therapies curbing net growth rates, but maintaining them at or above zero, could be interpreted as resulting from the action of either cytotoxic and/or cytostatic processes. In contrast, therapies reducing net growth rates below zero necessarily have a cytostatic component. Our model, or modifications of it to explicitly include cytotoxic and cytostatic effects, could be used in future research to make predictions about optimal dose and start times to achieve acceptable levels of tumor control, or the probability of a given tumor size by a given age.

Decisions whether or not to employ specific chemopreventive therapies carry with them the risk of a poorer outcome than would have been the case had another available strategy, or no treatment at all, been adopted [47]. This issue is relevant to situations where alterations in life-style, removal or treatment of pre-cancerous lesions, or medications potentially result in unwanted side effects or potentially induce new invasive neoplasms (e.g., [48]). Chemopreventive management prior to clinical detection would be most appropriate for individuals with genetic predispositions, familial histories, elevated levels of specific biomarkers, or risk-associated behaviors or life-styles [15,17,18, 49, 50]. Importantly, our approach presupposes that the danger a nascent, growing tumor presents is proportional to its size and (implicitly, all else being equal) a person’s age. Due caution is necessary in applying our results, since studies have argued that metastatic potential rather than tumor size may be a better predictor of future survival [51–53].

### Reactive approaches

Over the past decade, several alternative approaches to MTD have been proposed, where the objective is to manage rather than eradicate tumors (e.g., [8,12–14,54,55]). Tumor management attempts to limit cancer growth, metastasis, and reduce the probability of obtaining resistance mutations through micro-environmental modification or through competition with non-resistant cancer cell populations or with healthy cells. These approaches usually involve clinically diagnosed cancers: either inoperable tumors or residual cancers after tumor excision. In the former situation tumors are typically large enough in size to contain numerous resistance mutations. In many, if not most, cases these neoplasms will have metastasized, meaning greater variability both in terms of phenotypes and hence potential resistance to chemotherapies, and in penetrance of therapeutic molecules to targeted tumor cells [56, 57]. The latter situation involves smaller, residual cancer cell populations, but composed of high frequencies of resistant variants or dormant cells [56]. According to our results, both scenarios are likely to involve populations with large numbers of accumulated driver mutations (or fewer driver mutations, but each with larger selective effect), which ostensibly contribute to the speed of relapse. Thus, management of clinically detected tumors need not only limit the proliferation and spread of refractory subpopulations (Figures 3–4), but should also aim to control the growth of multi-driver subclones (Figure 4**B**).

We suggest that the frequency distribution of driver mutations and the distribution of resistant subclones within a heterogeneous cancer cell population could be used to instruct decisions of the time course of treatment levels, with the aims of curbing tumor growth, metastasis, and resistance. We found that tumors typically achieve several additional driver mutations by the time they reach detection (Figure 5**A**,**C**; Supplementary figure 3**A**, **C**), which approximates certain estimates [58], but falls short of others [59].

In conclusion, our results indicate that the single most important variable in determining therapeutic outcome is the size of the initial cancer cell population (i.e. when prevention commences and/or following resection). This highlights the importance of biomarkers as accurate proxies of otherwise undetectable malignancies [60], and the accurate assessment of micro-metastases [61]. We suggest that if order-of-magnitude estimates are possible, then low dose, continuous, constant approaches could be optimized. According to our model such options will always be superior to more aggressive chemotherapies.

## Acknowledgments

The authors are grateful to Athena Aktipis, Daniel Fisher, Sylvain Gandon, Urszula Hibner, Patrice Lassus, Carlo Maley and Ville Mustonen for discussions and helpful remarks. All calculations were made using programs, written in C, and the free, open-source statistical package R [63]. The color palette for figures was adopted from [64]. Code for all calculations, and for producing all of the figures, is available at [65] and can be used freely for non-commercial purposes. The work was made possible by the facilities of the Shared Hierarchical Academic Research Computing Network (SHARCNET:www.sharcnet.ca) and Compute/Calcul Canada. ARA was supported by CNRS Interdisciplinary postdoctoral program. MEH thanks INSERM “Physique Cancer” (CanEvolve PC201306), ANR (EvoCan ANR-13-BSV7-0003-01) and PICS (PlCS05313) for financial support.

## Supplementary figures

**Supplementary figure 1.**
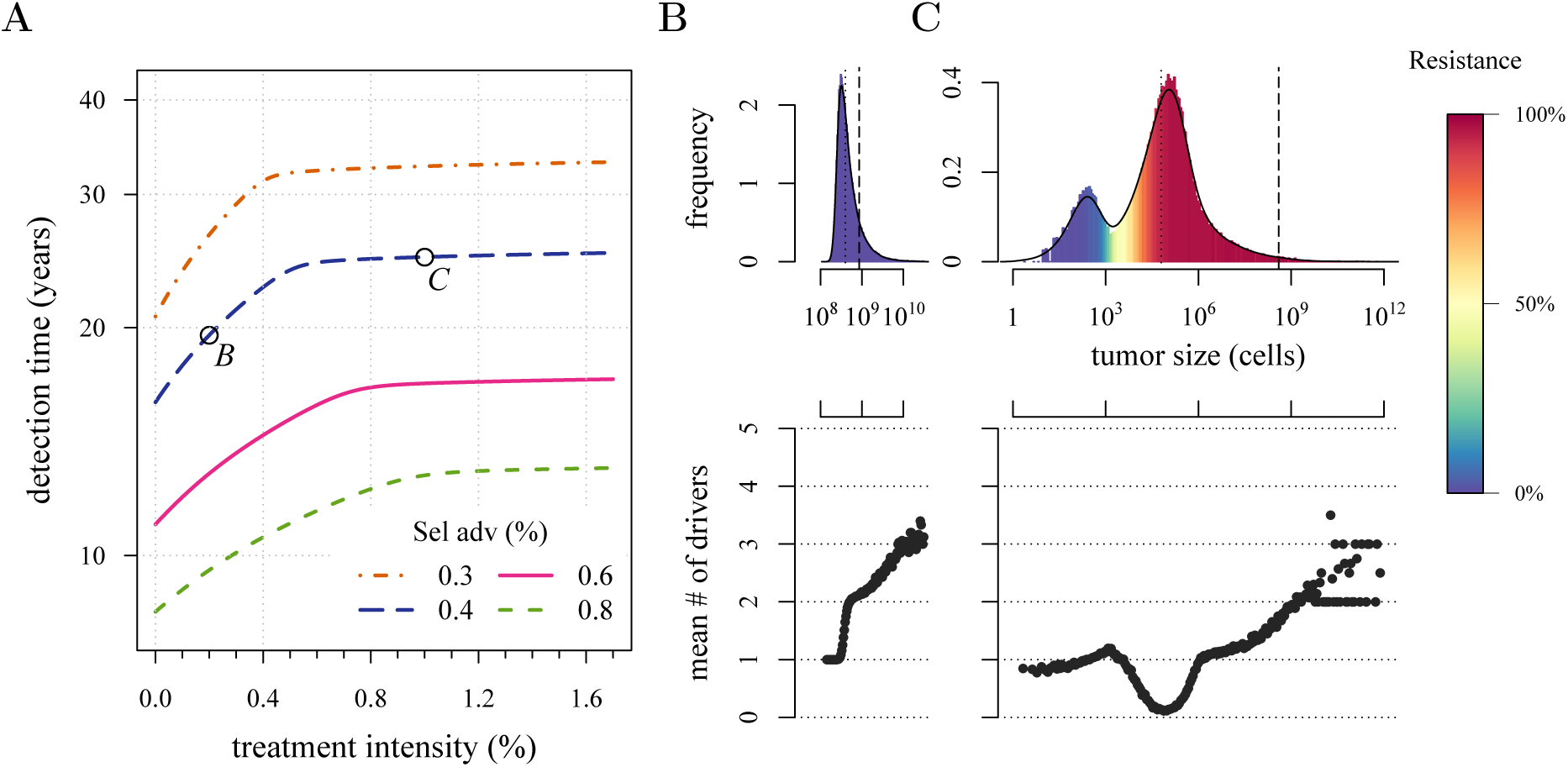
Tradeoff between growth and resistance under different treatment regimes. (**A**) Analytically-derived times for a tumor to reach 10^9^ cells (see equation (S11)). (**B**) and (**C**) Sample distributions for corresponding points *B* and *C*, shown in plot **A**. The bottom panel shows the mean number of additionally accumulated drivers for all detected tumors over intervals of 3 months. The color-code indicates the level of resistance in detected tumors over these intervals. Parameters otherwise as in Table 1.

**Supplementary figure 2.**
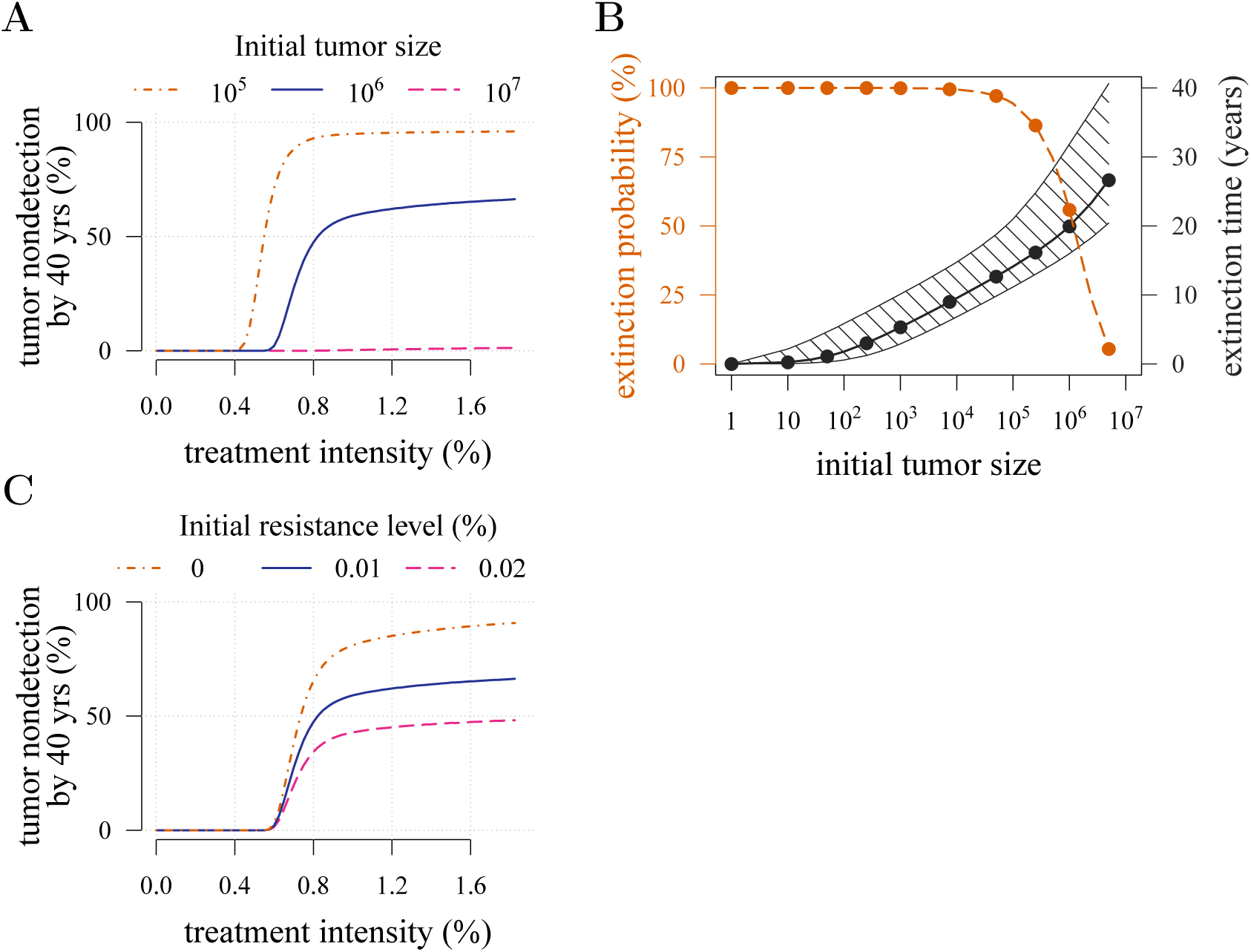
Effects of initial neoplasm size. (**A**, **B**) and resistance level (**C**) on preventive measure success. (**B**) The median (black line) and 90% confidence intervals (hatched area) for the distribution of extinction times. Red dashed line indicates the probability of tumor extinction, depending on initial cell number. Treatment level is 1.0% per cell cycle, and we assume no pre-resistance. Other parameters as in Table 1.

**Supplementary figure 3.**
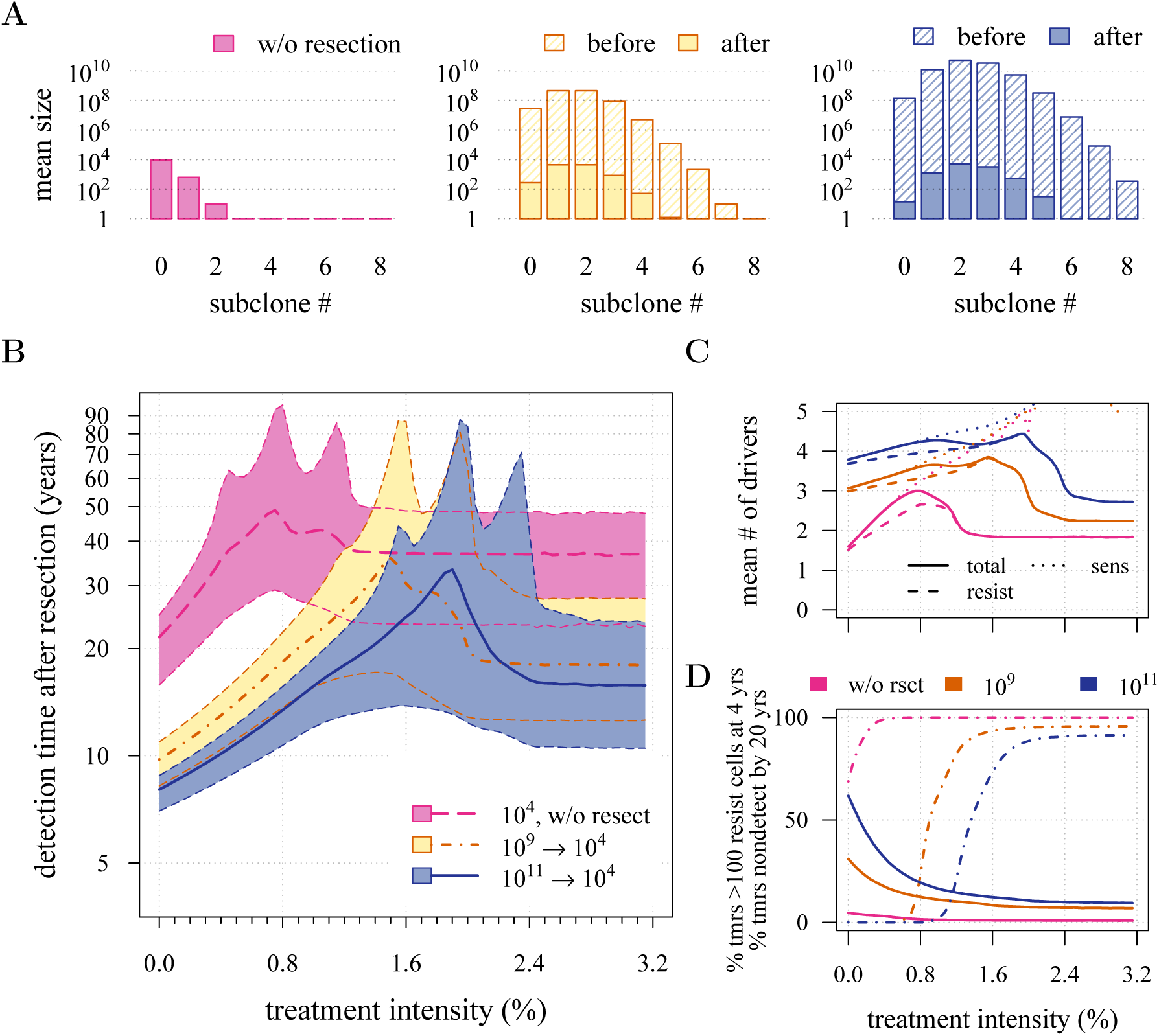
Preventive versus reactive measures. Therapeutic outcomes of prevention starting at 10^4^ cancer cells, versus resection leaving 10^4^ cells. (**A**) The distribution of mean sizes of subclones for different constant treatment intensities (hatched bars = before removal and solid bars = post removal). (**B**) The distribution of detection times (thick lines = medians, shaded areas with dashed boundaries = 90% CIs). (**C**) The mean number of accumulated drivers within a tumor at the time of detection. (**D**) The percentage of cases where the tumor consists of less than 100 resistant cells at 4 years after treatment commences (solid lines), and the percentage of cases where tumor size is below the detection threshold 20 years after the measure begins (dashed-and-dotted lines). Maximal number of additional drivers is 9, other parameter values as in Table 1.

**Supplementary figure 4.**
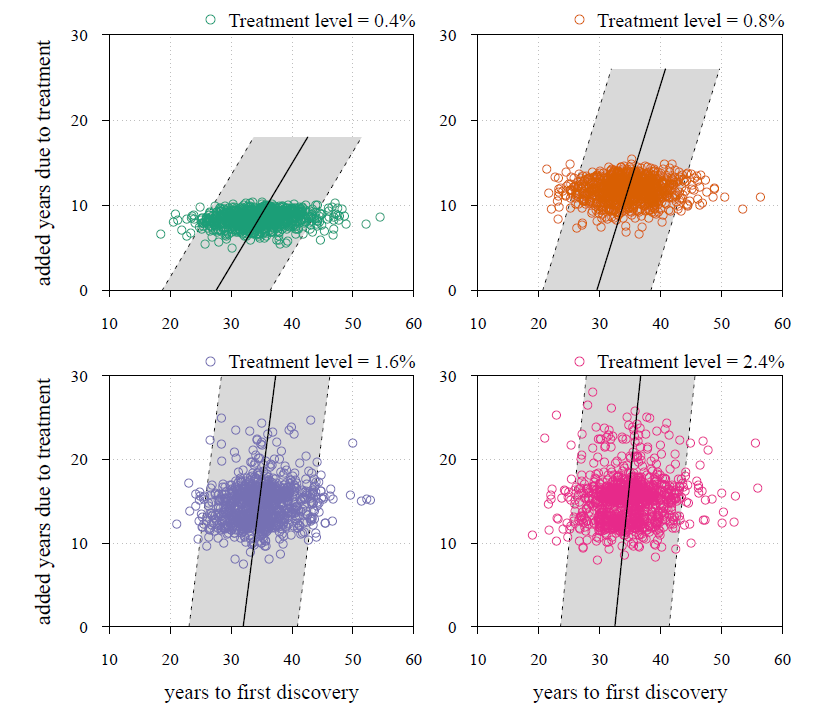
Time to first discovery as a predictor of treatment success. Time to tumor relapse following resection as function of the time it takes for the initial cancer cell to attain 10^9^ cells (i.e., the point at which the tumor is discovered, resected and treatment begins). Each dot represents a numerical simulation from the yellow distribution in Figure 5**B**. Four different treatment levels are considered. Black solid line is a simple linear regression and grey area with dashed boundaries indicates extrapolation of high and low bounds accounting for 95% of observations (prediction interval). Maximal number of additional drivers is 9. Other parameter values as in Table 1.

**Supplementary figure 5.**
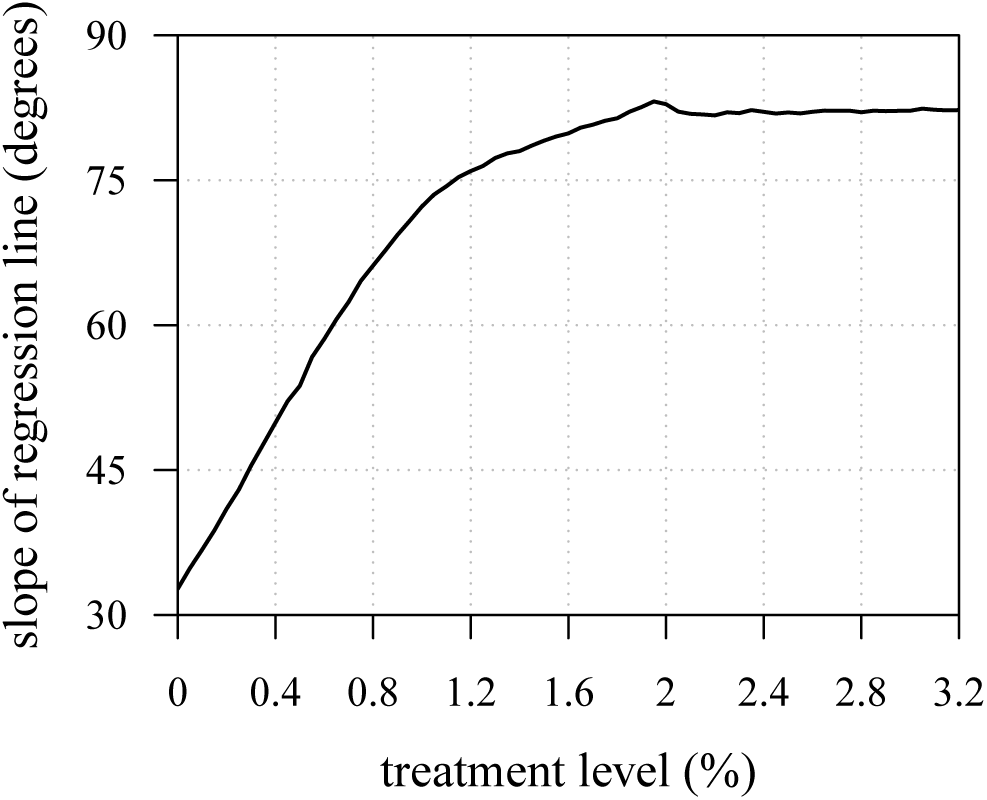
Time to tumor discovery is predictive of therapeutic outcome for low treatment levels. The slopes of regressions from numerical experiments for different treatment levels of time to tumor relapse following resection as function of the time it takes for the initial cancer cell to attain 10^9^ cells. See Supplementary figure 4 for details.

**Supplementary figure 6.**
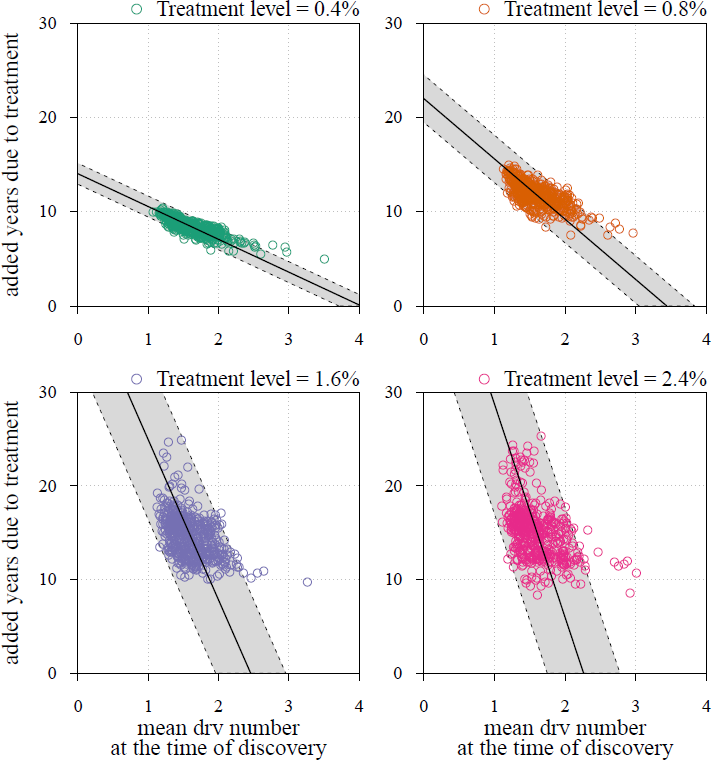
Mean number of drivers in resected tumor as a predictor of treatment success. See Supplementary figure 4 for other details.

**Supplementary figure 7.**
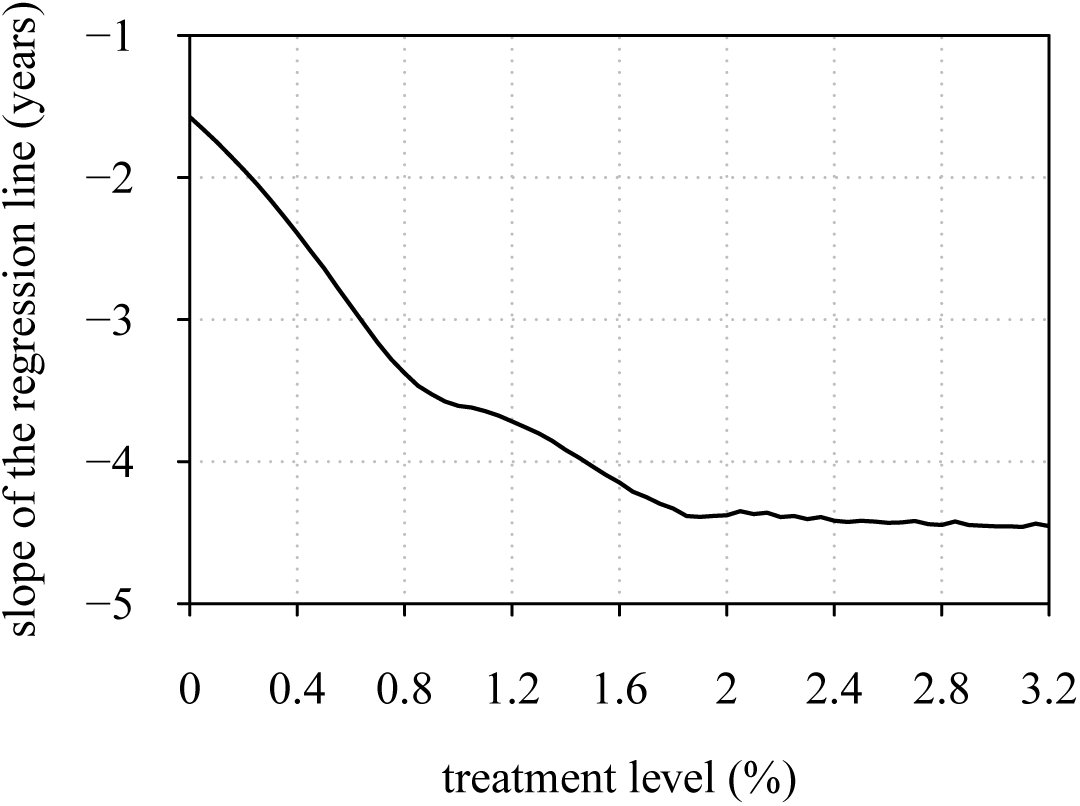
Time to tumor discovery is predictive of therapeutic outcome for low treatment levels. The slopes of regressions from numerical experiments for different treatment levels of time to tumor relapse following resection as function of the mean number of drivers in a resected tumor. See Supplementary figure 4 for details.

## Supplementary videos

**Supplementary videos 1–3.** Please visit: http://tiny.cc/AkhmHoch15VideoS123.

**Treatment level affects both detection time and frequency of resistance.** (**A**) The median (thick line) and 90% confidence intervals (shaded areas with dashed boundaries) for the distribution of detection times. Parameter values are as in Table 1 except the one being varied (see additional information before each video). (**B**) Particular samples of the distribution of detection times and distribution of the mean number of accumulated drivers. Color-code indicates the level of resistance in detected tumors over 3 month intervals.

**Supplementary videos 4, 5.** Please visit: http://tiny.cc/AkhmHoch15VideoS45.

**Comparison of preventive and reactive strategies.** (**A**) The median (thick line) and 90% confidence intervals (shaded areas with dashed boundaries) for the distribution of detection times. Parameter values are as in Table 1. (**B**, **C**) Particular samples of the distribution of detection times for preventive and reactive treatments respectively. The rectangle on the top of **B** or on the bottom of **C** shows the 5^th^ and 95^th^ percentiles, the blue circle indicates the median, while the red line - the mean of the distribution of detection times. Color-code indicates the mean number of additionally accumulated drivers for a period of 1 year.

## Supplementary information

### Mean-field approach

We use the mean-field approach, see e.g. [30], which approximates the behavior of a system consisting of many cells, so that the effects of stochasticity are averaged and an intermediate state is described by a set of ordinary differential equations.

### Master equations

We write master equations to track the probability *P_ij_*(*t*) that a randomly chosen cell from a population of tumor cells is of type (*i, j*) at time *t*.

The temporal dynamics of probabilities *P_ij_*(*t*), *i* = 0, 1,…, *N*, where *N* is the maximal number of additionally acquired drivers and *j* = 0, 1, are described by

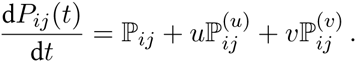

Here, the right-hand side is a superposition of probabilistic in- and out-flows from different mutational states to the current one (*i, j*). The function 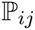 describes the growth of subclone (*i, j*) and is proportional to the probability *P_ij_*(*t*), multiplied by the difference between fitness *f_ij_* and its average value over the whole population 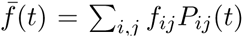. Functions 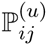 and 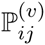 represent the probabilistic flows of mutations. For 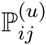, a driver is added from class (*i* − 1, *j*) to (*i, j*) in proportion to the probability *P_i_*_−1,*j*_(*t*), the probability of cell birth *b_i_*_−1,*j*_, and the probability of a zero locus being chosen from *N* total loci consisting of (*N* − (*i* − 1)) other zero loci. A similar approach is used to define the outflow term for the probability from class (*i, j*) to (*i* + 1, *j*). The second term 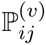 is the probability of mutating to therapeutic resistance (*i, j* = 0) to (*i, j* = 1), and is proportional to *P_i_*_0_(*t*) and birth rate *b_i_*_0_. Finally, all terms are summed, taking into account the initial conditions: *P*_00_(0) = 1 − *κ*, *P*_01_(0) = *κ* and *P_ij_* = 0 for any other *i* or *j*.

The above elements lead to the following system of ordinary differential equations (ODEs):

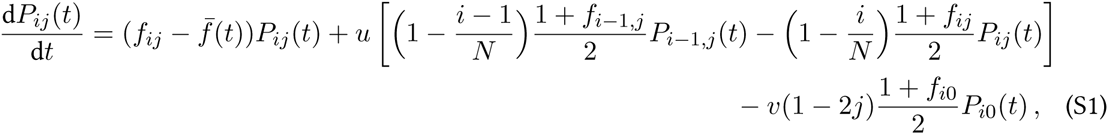

where some probabilities *P_ij_*(*t*) could, theoretically, take on negative values, e.g. *P*_−1*,j*_(*t*), when *i* = 0, in which case, they are set to zero.

A simple transformation

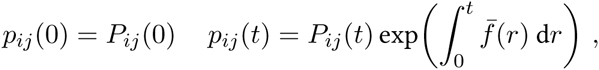

allows omitting the term 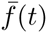 from Eq. (S1) and to linearize the latter with respect to the new “transformed” probabilities *p_ij_*(*t*). This gives

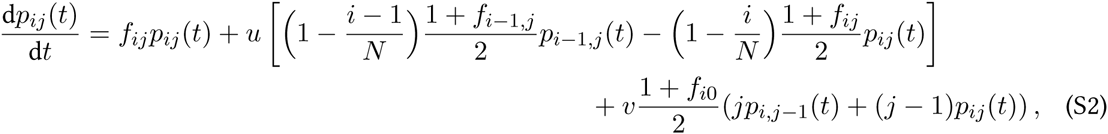

where, for convenience, we write (*jp_i,j_*_−1_(*t*) + (*j* − 1)*p_ij_*(*t*)) instead of (1 − 2*j*)*p_i_*_0_(*t*) (*j* = 0, 1).

### Probability generating function approach

With the master equations (S2), we apply the probability generating function (p.g.f.) method (31,66) to transform the system of (2*N* + 1) ODEs to a Hamilton-Jacobi (HJ) equation, that is, a first order partial differential equation.

We define the p.g.f. as the polynomial over all modified probabilities *p_ij_*(*t*) of the form

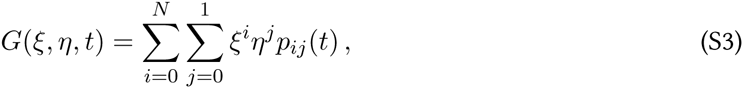

where *ξ* and *η* are variables that can be viewed as the momentum of an auxiliary Hamiltonian system governing the leading-order stochastic dynamics of the system [67]. Notice that the function *G*(*ξ*, *η*, *t*) is linear with respect to *η*.

Suppose that the function *G*(*ξ*, *η*, *t*) is defined, one can then obtain all characteristics of the stochastic process such as the average tumor size *n*(*t*) and the average frequency *n_res_* (*t*)/*n*(*t*) of resistant cells within a tumor. The former quantity is

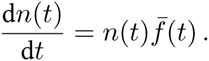

Using the normalization condition for the probability: 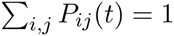, we obtain

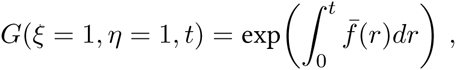

and then

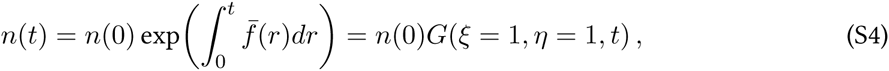

where the initial tumor size *n*(0) is sufficiently large. The frequency of resistant cells is defined as follows

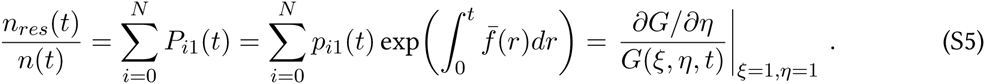

Initial conditions yield *p*_00_(0) = 1 − *κ*, *p*_01_ (0) = *κ* and *p_ij_* (0) = 0 for any other *i* and *j*, so that *G*(*ξ*, *η, t* = 0) = 1 − *κ* + *κη*.

To obtain the HJ equation related to the p.g.f. *G*(*ξ, η, t*), we multiply (S2) on *ξ^i^η^j^* and sum up all equations for *i* = 0, 1,…, *N* and *j* = 0, 1. After some algebra, we obtain

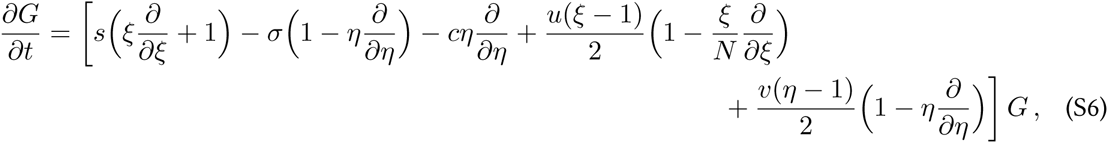

where only terms of order greater than or equal to *u*, *v* are retained, meaning that terms composed of the products *s*, *c* and *u*, *v* are omitted.

Equation (S6) is solved by the method of characteristics such that the HJ equation is transformed into a system of ordinary differential equations (i.e., the system of characteristics, see e.g. [68]).

### Time-varied treatment schedule

We find the characteristics for the variables *ξ* and *η* using (S6):

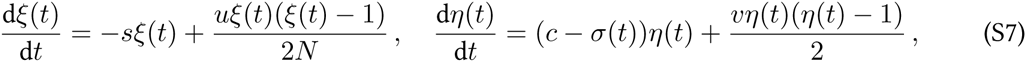

where *σ*(*t*) is a given function of time.

The p.g.f. *G*(*ξ, η, t*) changes along the characteristic (S7) according to the following ODE

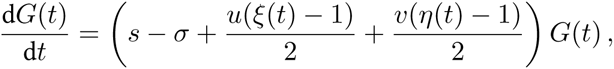

which is straightforward to integrate. Indeed, if we use (S7), this yields: d ln *G* = (*s* (*N* + 1) − *c*)d*t* + *N*d ln *ξ* + d ln *η*. Thus,

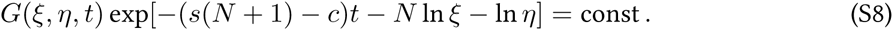

Recall that the quantity on the left hand side remains constant only along the characteristic curve (S7).

To obtain *G*(*ξ* = 1, *η* = 1, *t*), we need to solve (S7) subject to *ξ*(*t*) = *η*(*t*) = 1 and find *ξ*(0) and *η*(0). Then, given the initial condition *G*(*ξ*(0), *η*(0), 0) = 1 − *κ* + *κη*(0), *κ* is a level of resistance within a tumor (*κ* ∊ [0,1]), and we can finally define *G*(*ξ, η, t*) using (S8).

Finally, we use (S4) to derive the dynamics of *n*(*t*). To obtain the mean frequency of resistant cells within a tumor, we first write *∂G*/*∂η*, using (S8) with the right hand side implicitly dependent on *η*. (Note that time *t* is measured in cell cycles, which are assumed to be of 4 days on average. To derive all necessary equations with respect to the actual time, we need to divide *t* by the length of the cell-cycle *T* and substitute it in the equations.)

### Constant treatment

We study the case for constant *σ*. Notice that this includes the case of no treatment (*σ* = 0).

First, we find the characteristics for the variables *ξ* and *η*. Namely, solution of (S7) gives

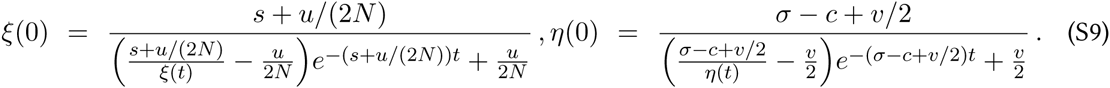

The subsequent substitution of (S9) into (S8) leads to

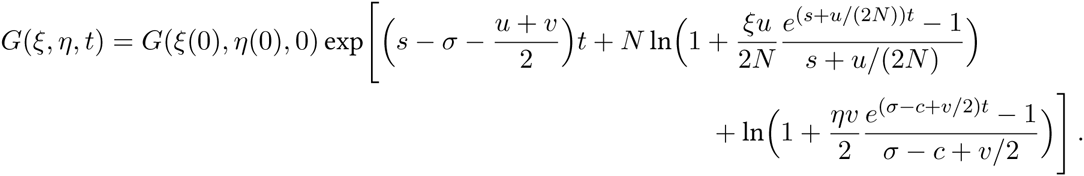

Taking into account *u, v* ≪ *s, c* and assuming *v* ≪ *σ* − *c*, we simplify further and write its approximate form

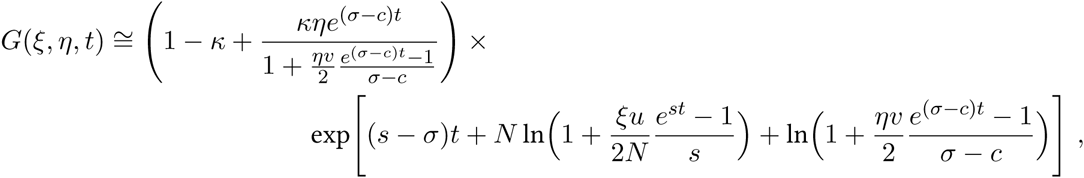

which can be also written in the form

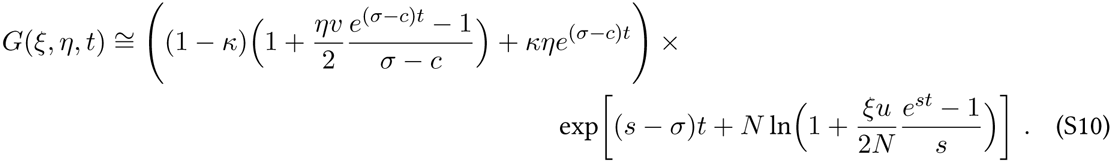

As expected (S10) is linear with respect to *η*.

Thus, we derive an analytical expression for the dynamics *n*(*t*). Namely, we use (S4), (S10) and substitute *ξ* = *η* = 1, to obtain

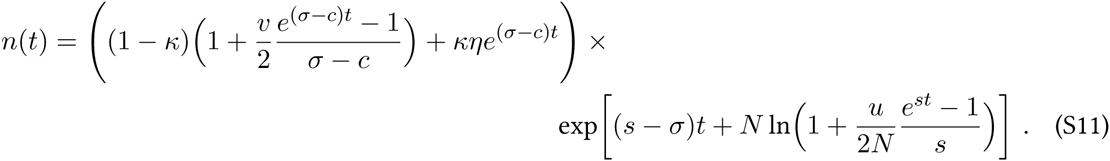

Equation (S11) is simplified for two limiting cases. In the early stages of tumor growth, the value *n*(*t*) changes according to a hyper-exponential law

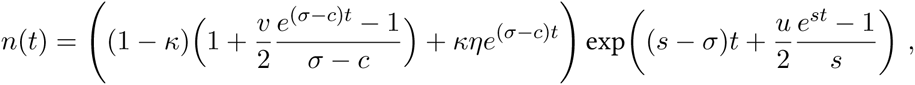

while at later stages the most aggressive subclone persists, being sensitive if *σ* < *c* (*n*(*t*) ∝ *e*^*s*(*N*+1)*t*^) and resistant otherwise (*n*(*t*) ∝ *e*^(*s*(*N* + 1)−*c*)*t*^).

To compute the frequency of resistant cells within a tumor (S5), we derive *∂G*/*∂η* using (S10):

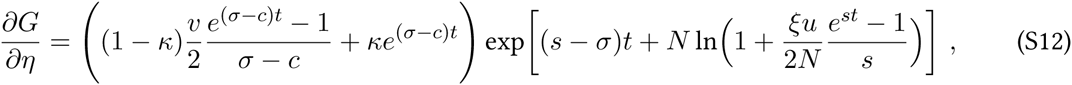

so that

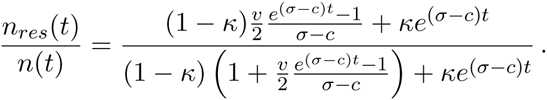

### Distribution of subclones within an exponentially growing tumor

The p.g.f. *G*(*ξ, η, t*) is used to derive expressions for all *P_ij_* (*t*), which are the probabilities of selecting a cell of type (*i, j*) from a tumor at time moment *t*. Namely, we need to differentiate the p.g.f. with respect to *ξ* and *η*, so that

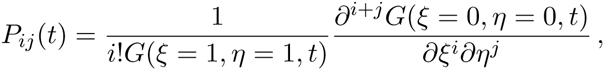

where *i* = 0, 1,…, *N* and *j* = 0, 1. Thus, we write

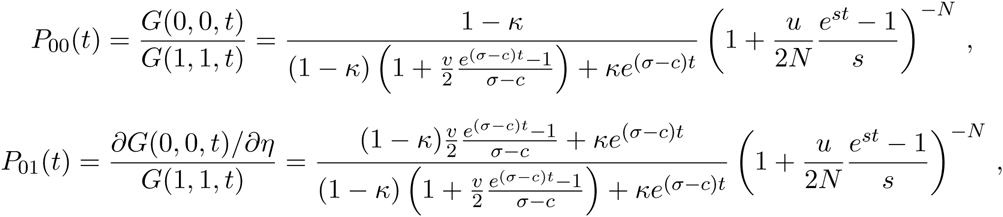

then

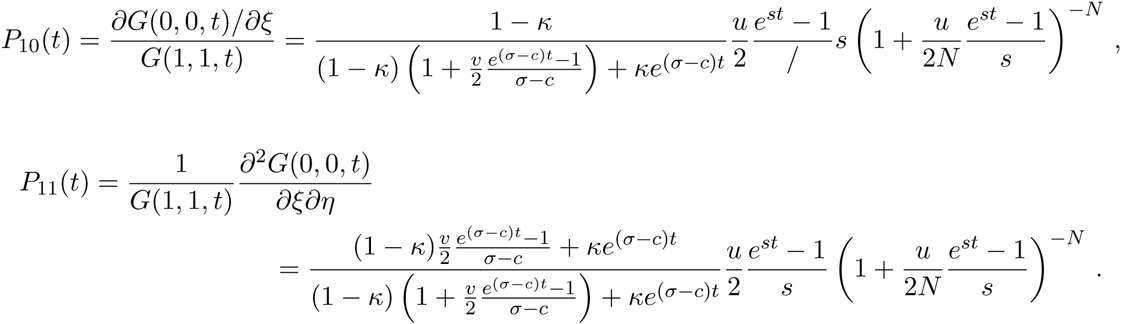

The general formula is written as follows

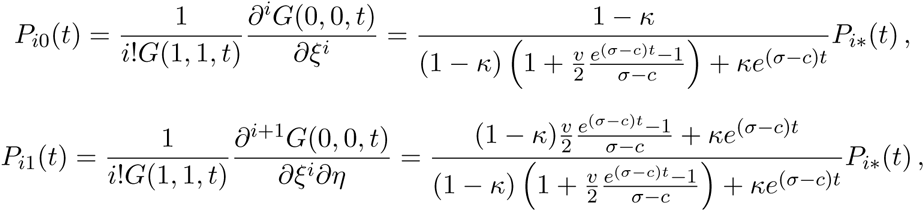

where *i* = 0, 1,…, *N* and the function

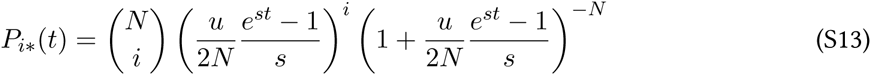

**Supplementary figure 8.**
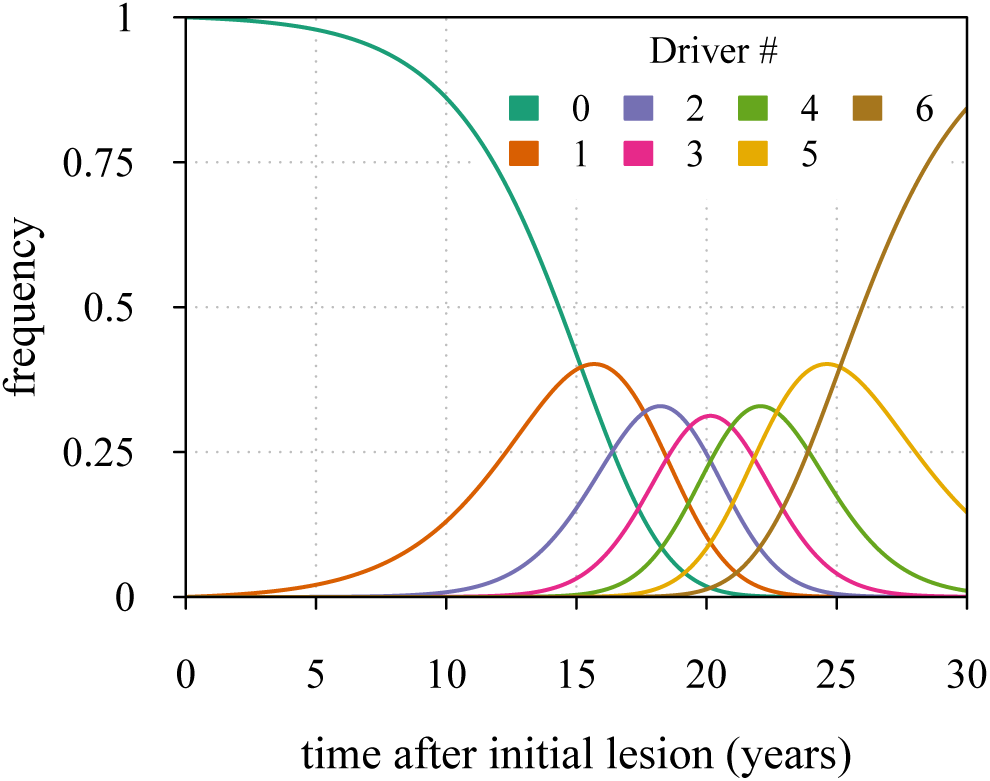
Maximal tumor heterogeneity in terms of driver subclones occurs at intermediate times after initial lesion.

defines the probability to pick a cell with *i* drivers independently of resistant status, 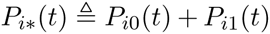, 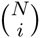 denotes a binomial coefficient, equal 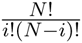.

The distribution *P_i*_*, (*t*) for a particular case of *N* = 6 is shown in Supplementary figure 8.

We now derive the mean time period when a given subclone with *i* additionally accumulated drivers dominates within a tumor.

Defining the time moments *t* = *t_i_* for which *P_i_*_−1*,\**_ (*t_i_*) = *P_i*_*(*t_i_*) (*i* = 0, 1, 2,…, *N*) gives

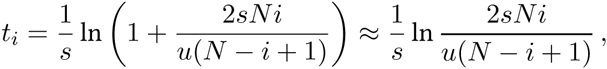

where we assume *u* ≪ *s*.

The time period when the subclone with *i* drivers prevails in a cell population is defined by the following expression

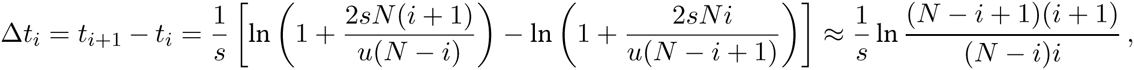

where *i* ≠ 0. For *i* = 0, we have

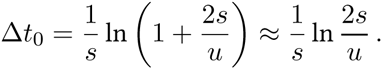

The latter formula has been previously reported (see Eq.(S7) in reference [27]).

